# MoCoO: Momentum Contrast ODE-Regularized VAE for Single-Cell Trajectory Inference and Representation Learning

**DOI:** 10.64898/2026.03.27.714791

**Authors:** Zeyu Fu

## Abstract

Characterising cellular differentiation from single-cell RNA sequencing (scRNA-seq) requires representations that capture both discrete cell-type identity and continuous developmental trajectories. We present MOCOO, a modular framework integrating a Variational Autoencoder (VAE), Neural Ordinary Differential Equations (Neural ODE), and Momentum Contrast (MoCo), complemented by a systematic Phase-2 Flow Matching (FM) refinement step applicable to all model variants. Through a systematic six-configuration ablation across 20 scRNA-seq datasets evaluated with a proposed five-metric suite covering clustering geometry (ASW, DAV, CAL) and embedding quality (DRE, DREX), we demonstrate two central findings. First, the ODE+MoCo combination is the core architectural synergy: VAE+ODE+MoCo achieves four of five top-two finishes among base configurations, including the best ASW (0.225) and DAV (1.478), plus second-best DRE (0.640) and CAL. Second, FM refinement systematically improves both embedding quality and clustering geometry across all six base configurations—DREX in 92% and DRE in 88% of 120 dataset– configuration pairs (ΔDREX= +0.030, ΔDRE= +0.023), CAL in 88%, ASW in 86% (ΔASW= +0.018), and DAV in 80% (ΔDAV= −0.072; Fig. 2). Combined, the full MoCoO pipeline (VAE+ODE+MoCo+Proto+FM) achieves the best DRE (0.678), DREX (0.660), and CAL, while VAE+ODE+MoCo+FM achieves the best ASW (0.257) and DAV (1.359). ODE smooths the latent manifold along developmental trajectories; MoCo sharpens cluster geometry; FM recovers and amplifies both embedding quality and cluster separation post-hoc. Downstream validation confirms that MoCoO latent spaces support annotation transfer, uncertainty quantification, differential expression, and branching detection. Pseudotime predictions correlate significantly with canonical marker genes across all five core developmental systems. We publicly release the MoCoO Python package (`pip install mocoo`) and full benchmark suite.

## I. Introduction

Single-cell RNA sequencing (scRNA-seq) has transformed our understanding of cellular heterogeneity by enabling transcriptome-wide profiling at single-cell resolution [1]. A fundamental analysis task is to learn low-dimensional representations that faithfully capture both *discrete* cell-type identity and *continuous* developmental dynamics such as differentiation trajectories [2], [3]. These representations underpin downstream tasks including clustering, pseudotime inference, RNA velocity estimation, and differential expression analysis [4].

Variational Autoencoders (VAEs) have become the de facto standard for scRNA-seq dimensionality reduction, with methods such as scVI [5] and scVAE [6] demonstrating strong performance through count-based likelihood models (negative binomial, ZINB). However, standard VAE latent spaces tend to be over-smoothed, conflating nearby cell states. Two complementary strategies address this limitation.

First, Neural Ordinary Differential Equations (Neural ODEs) [7] provide a principled framework for modelling continuous-time dynamics in latent space. By parameterising the latent derivative with a neural network and integrating via an ODE solver, these models learn smooth trajectories without discrete time-step assumptions. Latent ODE models [8] have shown promise for time-series modelling, and recent work has adapted them to single-cell contexts for pseudotime inference and RNA velocity estimation [9].

Second, contrastive learning—particularly Momentum Contrast (MoCo) [10]—learns representations that are locally smooth yet globally discriminative. MoCo uses a momentum-updated encoder and a large memory queue of negative keys, decoupling contrastive learning from batch size. Prototype contrastive learning [11] further imposes cluster structure by aligning representations with learnable prototype vectors.

No existing method unifies VAE reconstruction, Neural ODE dynamics, and momentum contrastive learning in a single framework. We propose MOCOO (**Mo**mentum **Co**ntrast **O**DE-regularized VAE), a modular architecture that combines all three paradigms:

**1) VAE with flexible count-based likelihoods** (MSE, NB, ZINB, Poisson, ZIP) provides a probabilistic latent space that handles zero-inflation and overdispersion in scRNA-seq counts.

**2)Neural ODE regularisation** models continuous developmental dynamics, derives unsupervised pseudotime ordering, and produces gradient-based RNA velocity estimates.

**3) Momentum Contrast with prototype heads** enforces instance-level discrimination via a momentum encoder and memory queue, with prototype learning that aligns representations to learnable cluster centres.

**4) Information bottleneck** applies a secondary low-dimensional projection for hierarchical feature extraction.

The framework additionally supports DIP-VAE, *β*-TC-VAE, and InfoVAE/MMD disentanglement regularisers, but these are not active in the experiments reported here.

We conduct a systematic ablation across six configurations (VAE, VAE+ODE, VAE+MoCo, VAE+MoCo+Proto, VAE+ODE+MoCo, and VAE+ODE+MoCo+Proto) on 20 scRNA-seq datasets spanning diverse developmental systems, with a Phase-2 Flow Matching (FM) refinement step that extends this to 12 configurations. Deep-learning external baselines with comparable latent representations—scVI [5], Harmony [12], and scANVI [13]—contextualise MoCoO’s absolute performance. A key finding is that combining ODE and MoCo consistently produces the best cluster geometry (cross-dataset mean ASW 0.225, DAV 1.478 for VAE+ODE+MoCo), while VAE+ODE achieves the best embedding quality (DRE 0.647, DREX 0.613). ODE-containing configurations consistently outperform non-ODE variants on cluster geometry across all 20 datasets (ΔASW ∼0.04, ΔDAV ∼0.23). FM refinement improves distance-preservation metrics in over 85% of dataset–configuration pairs (DREX Δ= + 0.030, DRE Δ= + 0.023). Component analysis reveals that each module contributes complementary geometric structure: the Neural ODE smooths the latent manifold along trajectories and MoCo sharpens pairwise distance relationships, with the ODE+MoCo combination producing the strongest overall synergy across all 20 datasets. Evaluation is centred on a proposed five-metric suite covering clustering geometry (ASW, Davies–Bouldin, Calinski–Harabasz) and embedding quality (DRE, DREX), complemented by biological validation (pseudotime–marker correlations), downstream analysis (annotation transfer, uncertainty quantification, generation quality, gene importance, branching detection, differential expression), batch integration (iLISI, bASW, cLISI, graph connectivity, isolated label ASW via scIB [14]), and training dynamics.

## II. Related Work

### A. Variational Autoencoders for scRNA-seq

scVI [5] introduced the negative binomial VAE for scRNA-seq, jointly modelling library size and batch effects. Extensions include scVAE [6] (multiple count distributions), scANVI [13] (semi-supervised cell-type annotation), and to-talVI [15] (multi-omics). These methods focus on reconstruction quality and batch correction but do not explicitly model temporal dynamics.

### B. Neural ODEs and Latent ODE Models

Neural ODEs [7] parameterise continuous dynamical systems 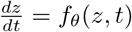 and integrate using adaptive ODE solvers. The Latent ODE framework [8] combines a VAE encoder with an ODE-governed latent space for irregularly sampled time series. Applications to single-cell biology include trajectory inference [9], while related dynamical modelling approaches have advanced RNA velocity estimation [16]. Optimal-transport-based methods such as TrajectoryNet [17] and CellOT [18] model population-level dynamics rather than single-cell ODE trajectories. MoCoO extends the PanODE framework [9] (by the first author). Specifically, MoCoO inherits the VAE encoder–decoder, Neural ODE integration, and pseudotime-prediction head from PanODE. The novel contributions of the current work are: (i) the MoCo module with momentum encoder and memory queue, (ii) learnable prototype heads for cluster anchoring, (iii) the information bottleneck, (iv) the systematic six-configuration ablation design, and (v) the DRE/DREX evaluation suites. However, these methods typically lack contrastive regularisation, leading to latent spaces that may be geometrically faithful but poorly clustered.

### C. Contrastive Learning for Single-Cell Data

MoCo [10] and SimCLR [19] have been adapted to biological contexts. scAGCL [20] applies symmetric augmentation-guided contrastive learning to scRNA-seq data with cell-type-aware positive pair construction. scGPCL [21] introduces prototype contrastive learning that aligns cell embeddings with learnable prototype vectors representing cell types. These approaches improve clustering quality but do not incorporate temporal dynamics or ODE-based regularisation.

### D. Integrated Approaches

Several methods combine two of the three paradigms. scDiff [22] uses diffusion models (related to score-based SDEs) for single-cell generation. VeloVAE [23] integrates VAE with RNA velocity. Monocle 3 [24] and PAGA [25] use graph abstractions for trajectory inference. For batch integration, Harmony [12] applies iterative soft clustering in PCA space to remove batch effects. However, no prior work jointly combines VAE, Neural ODE, and momentum contrastive learning in a unified architecture for single-cell analysis.

## III. Method

### A. Overview

MoCoO (Fig. 1) processes scRNA-seq count data *X* ∈ ℝ *N* ×*G* (*N* cells, *G* genes) through five components: (1) a VAE with count-based decoder, (2) a Neural ODE for continuous dynamics, (3) a Momentum Contrast module with memory queue, (4) an information bottleneck, and (5) optional disentanglement regularisers.

**Fig. 1.**
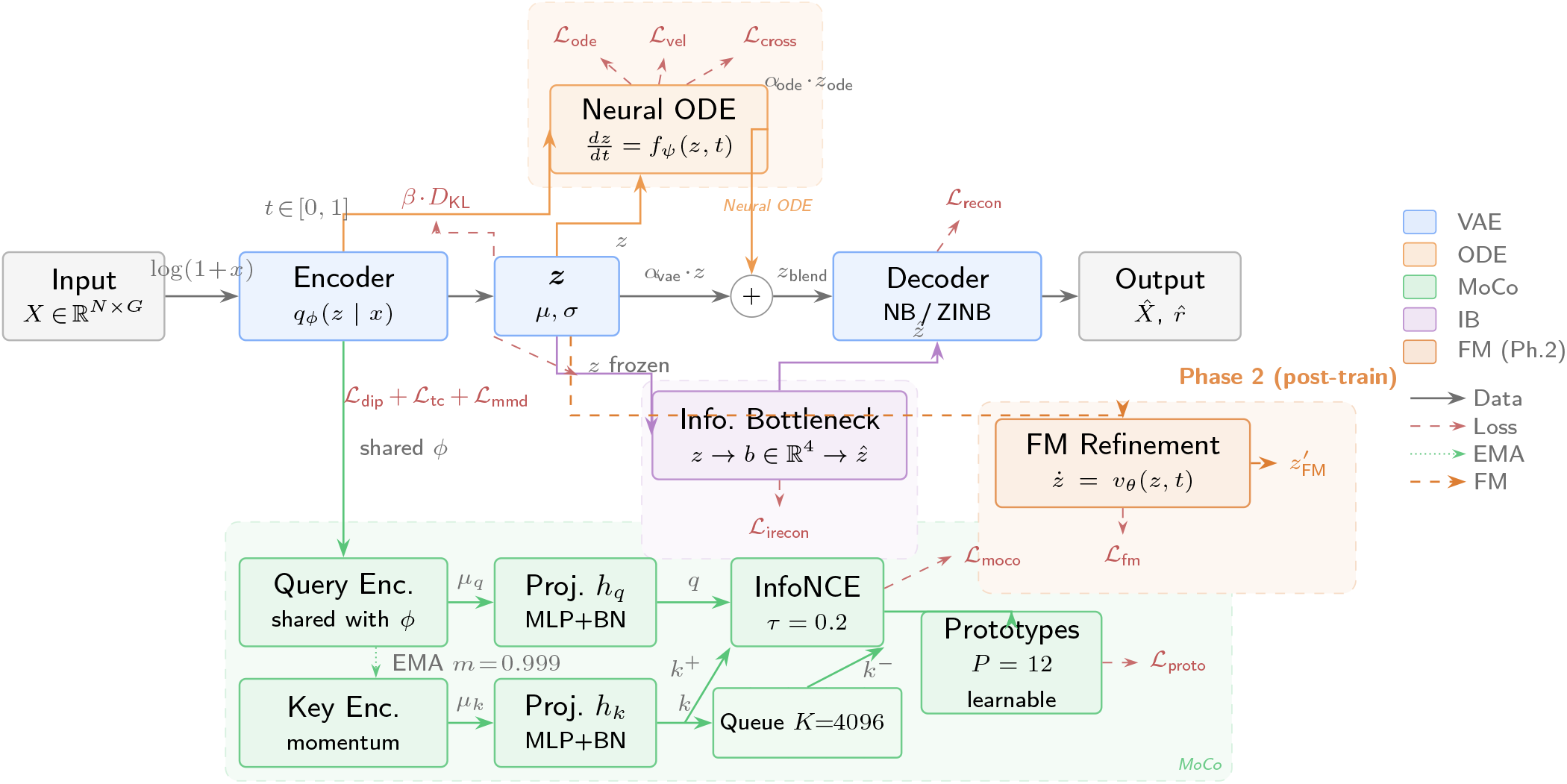
MOCOO architecture overview (**Fig. 1**). Count data *X* ∈ *ℝN×G* passes through a VAE encoder to produce latent representations *z*, refined by a Neural ODE solver (orange) for continuous trajectory modelling. Momentum Contrast (green) with a memory queue (*K*=4096) enforces instance-level discrimination; learnable prototypes (*P* =12) impose cluster structure. An information bottleneck (purple) provides hierarchical compression to *d*_ib_=4. Dashed regions denote modular components. Phase-2 FM refinement (vermilion, right) operates post-training on the frozen latent space of any configuration—the central contribution of this work. Colour coding is consistent throughout all figures.

### B. Encoder

The encoder *q*_*ϕ*_(*z*|*x*) maps log-normalised expression 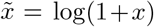 through a two-layer ReLU MLP to produce mean and pre-softplus scale *σ*^′^ of a diagonal Gaussian posterior (1); latent samples are drawn via reparameterisation (2):

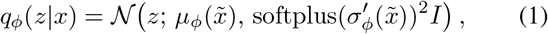

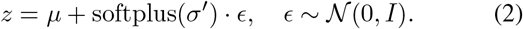

When ODE mode is enabled, the encoder also predicts a scalar pseudotime *t* ∈ [0, 1] via a sigmoid-activated linear head.

### C. Neural ODE Component

Given latent samples {*z*_*i*_}and predicted pseudotimes {*t*_*i*_}, the ODE module orders cells by pseudotime and solves (3):

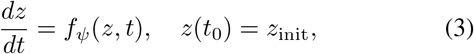

where *f*_*ψ*_ is a two-layer ELU MLP. Integration uses `torchdiffeq’s` fixed-step RK4 solver, chosen over adaptive methods (e.g., Dormand–Prince [26]) because the encoder-predicted pseudotime grid is relatively smooth and fixed-step integration provides deterministic gradients without adaptive step-size overhead—important for stable joint optimisation of the encoder and ODE function. Preliminary comparisons showed negligible clustering geometry differences (ΔASW< 0.005, ΔDAV< 0.02) between RK4 and Dormand–Prince adaptive integration, with ∼30% longer wall-clock time for the adaptive solver due to step-size adaptation overhead in the joint training loop. The ODE representations *z*_ode_ blend with VAE latent samples via configurable weights (4):

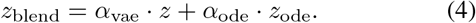

Two auxiliary losses regularise the ODE:

- **ODE–VAE alignment:**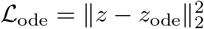
- **Velocity consistency: ℒ**_vel_ = 1− cos(*z*_*i*+1_−*z*_*i*_, *f*_*ψ*_(*z*_*i*_, *t*_*i*_)), aligning the learned velocity field with empirical displacements.

### D. Momentum Contrast Module

The MoCo module follows MoCo v1 [10] with enhancements from scAGCL [20] and scGPCL [21]:

1. **Query and Key Encoders:** The query encoder shares parameters with the VAE encoder; the key encoder is updated via EMA: *θ*_*k*_← *m*· *θ*_*k*_ +(1− *m*) ·*θ*_*q*_, *m* = 0.999.
2. **Projection Heads:** Two-layer MLPs (BatchNorm, ReLU) map *d*-dimensional latent vectors to a *d*_proj_-dimensional contrastive space.
3. **Memory Queue:** A FIFO buffer of size *K* stores projected keys, decoupling the number of negatives from batch size.
4. **InfoNCE Loss** (see (5)):

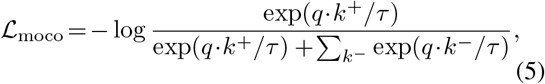

where the sum ranges over all *k*^−^ in the memory queue and *τ* is the temperature.
5. **Prototype Contrastive Loss** (optional; see (6)):

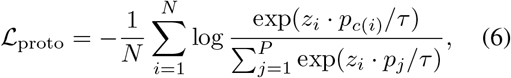

where 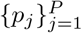 are learnable prototype vectors and *c*(*i*) = arg max_*j*_ *z*_*i*_ *p*_*j*_*/τ* assigns cell *i* to its nearest prototype.

#### Positive pair construction

For each mini-batch, two stochastic augmented views of the same cell form positive pairs: the query view passes through the main encoder, while the key view passes through the momentum encoder. The augmentation pipeline applies three transformations, each with probability *p*_aug_=0.5: (1) gene masking—randomly zeroing out genes with probability 0.1; (2) Gaussian noise—adding 𝒩 (0, 0.04) to randomly selected genes (probability 0.1); (3) feature swapping—exchanging gene values across cells in the batch (probability 0.05). Negatives come from all keys stored in the memory queue.

##### Augmentation rationale

Gene masking simulates dropout, which is a dominant noise source in scRNA-seq protocols [27]. Gaussian noise (*σ*=0.2) models technical variability in library preparation. Feature swapping operates at very low probability (*p*=0.05, affecting ∼5% of genes per cell), which is equivalent to a regularised mixup—it encourages invariance to minor expression perturbations without creating biologically implausible hybrid profiles. All three augmentations are deliberately conservative: masking affects only 10% of genes per view, noise is applied independently per gene, and swapping probability is below the natural inter-cell variance within a cell type. This design follows the principles validated in scAGCL [20] and scGPCL [21], which demonstrated that moderate stochastic augmentations improve contrastive learning for scRNA-seq without distorting cell-state semantics.

### E. Decoder

The decoder *p*_*θ*_(*x*|*z*) reconstructs count data from the latent representation. It predicts normalised mean parameters *ρ* = softmax(MLP(*z*)) and uses gene-specific learnable dispersions *r*:

- **Negative Binomial (NB):** *p*(*x*_*g*_|*z*) = NB(*x*_*g*_; *µ*_*g*_ = *l* · *ρ*_*g*_, *r*_*g*_)
- **Zero-Inflated NB (ZINB):** *p*(*x*_*g*_|*z*) = *π*_*g*_ · *δ*_0_(*x*_*g*_) + (1 − *π*_*g*_) · NB(*x*_*g*_; *µ*_*g*_, *r*_*g*_)

where *l* = Σ*_g_x*_*g*_ is the library size and *π*_*g*_ is a dropout probability predicted by a secondary decoder head. While MoCoO supports MSE, NB, ZINB, Poisson, and ZIP likelihoods, all experiments in this paper use the negative binomial (NB) likelihood unless otherwise stated.

### F. Information Bottleneck

A secondary encoder-decoder pair maps *z*∈ ℝ ^*d*^,*/sup>*to a bottleneck *b* ∈ 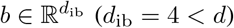and back to 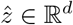, imposing hierarchical compression that retains only the most salient features. The bottleneck reconstruction loss 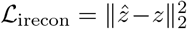is included in the total loss (8) with weight *λ*_irecon_ = 1.0. The IB serves as a fixed architectural component that forces the model to identify a low-dimensional summary of the latent representation; it is not ablated independently because preliminary experiments showed its effect to be minor compared to ODE and MoCo (clustering geometry changes < 0.005 ASW, < 0.02 DAV), and it adds negligible computational cost.

### G. Flow Matching Refinement

As a systematic post-training enhancement, MoCoO includes a *Phase-2 Flow Matching (FM)* refinement step that operates on the frozen latent space of any base configuration. Given the trained VAE encoder, FM learns a continuous normalising flow that transports a standard Gaussian prior *z*_0_∼𝒩 (0, *I*) toward the empirical latent distribution *z*_1_ = *µ*_*ϕ*_(*x*) via linear interpolation paths [28]: *z*_*t*_ = (1−*t*)*z*_0_ + *tz*_1_. A two-layer MLP velocity network *v*_*θ*_(*z*_*t*_, *t*) is trained to predict the target velocity *z*_1_ − *z*_0_ at each interpolation time *t* ∈ [0, 1] using the conditional flow matching loss:

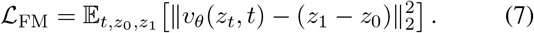

At inference, FM-refined latent codes are obtained by integrating *v*_*θ*_ from *t*_start_ to *t*=1 using Euler steps. Setting *t*_start_ < 1 blends learned flow dynamics with the encoder output; *t*_start_ = 1 recovers the original encoder latent (no FM). The FM step adds six base configurations as FM-augmented variants (e.g., VAE+FM, Full+FM), yielding 12 total configurations. Default settings: *t*_start_=0.9, FM epochs=200, lr=10^−3^, hidden dim=128, Euler steps=100.

### H. Total Loss

The total objective combines all active components (8):

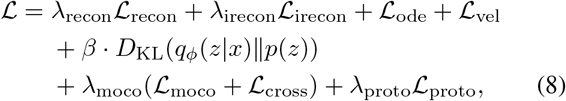

where each *λ* is a configurable weight. The cross-path contrastive loss 𝒩_cross_ aligns the ODE and VAE representations of the *same cell* via symmetric NT-Xent (9):

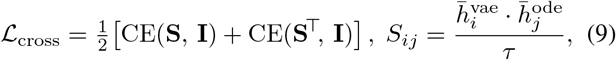

where 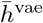 *ℓ*_2_-norm(proj(*z*)),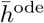 *ℓ*_2_-norm(proj(*z*_ode_)), **I** is the identity permutation (diagonal entries are positives), and CE denotes row-wise cross-entropy. The gradient of _cross_ flows only through the ODE path (*z* is detached), so the ODE aligns to the encoder rather than vice versa.

## IV. Experiments

### A. Datasets

We evaluate MoCoO on 20 scRNA-seq datasets spanning diverse biological contexts. Five core datasets are used for the main ablation study and downstream validation; an additional 15 datasets evaluate cross-dataset generalization and FM enhancement.

#### Core datasets (5)

1. **IRALL (Hematopoiesis):** 41,252 cells, 12 types, 8 batches (d0–d30). Mouse bone marrow HSC-to-mature-lineage differentiation.
2. **Dentate (Neurogenesis):** 18,213 cells, 14 types. Mouse dentate gyrus from radial glia to mature granule neurons.
3. **Endo (Endocrine Pancreas):** 2,531 cells, 13 types. Mouse pancreatic ductal-to-endocrine differentiation.
4. **Paul (Myeloid/Erythroid):** 2,730 cells, 19 types. Myeloid/erythroid bifurcation [29]; fine-grained with 19 clusters (some < 30 cells).
5. **Spinoids (Spinal Cord Organoid):** 9,619 cells, 8 types. Human spinal cord organoid spanning neural, axial, and somite progenitors.

#### Extended datasets (15)

Astrocyte, Brain Metastasis, Breast, Gastric, Hemato, Hepatoblastoma, hESC-time, Liver Cancer, Lung, Melanoma, Pituitary, Retina, Setty, Spine, and Teeth. These cover additional human and mouse tissues and developmental systems, providing a comprehensive generalization benchmark. All 20 datasets are processed with identical hyperparameters (tuned on IRALL only) and evaluated using all 12 configurations (6 base + 6 FM), yielding 20×12 = 240 configuration–dataset evaluation points. For external benchmarking (Section VI-E), 7 core developmental datasets (IRALL, Paul, Endo, Dentate, Hemato, Lung, Spinoids) are evaluated across all 18 baselines on all five proposed metrics.

### B. Model Configurations

Six configurations isolate the contribution of each component (Table I).

**TABLE 1.**
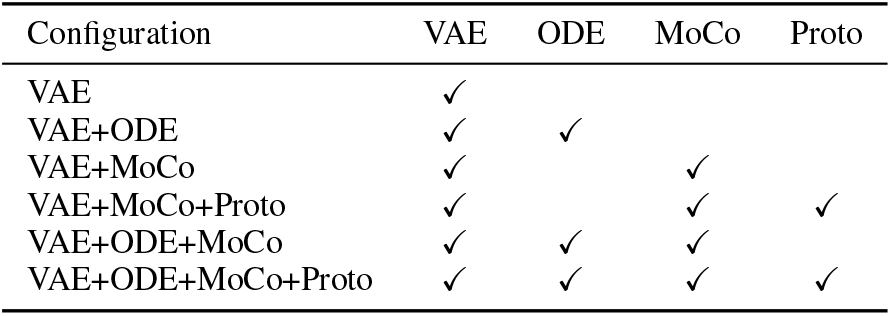
Model configurations and their active components.

| Configuration | VAE | ODE | MoCo | Proto |
| --- | --- | --- | --- | --- |
| VAE | ✓ |  |  |  |
| VAE+ODE | ✓ | ✓ |  |  |
| VAE+MoCo | ✓ |  | ✓ |  |
| VAE+MoCo+Proto | ✓ |  | ✓ | ✓ |
| VAE+ODE+MoCo | ✓ | ✓ | ✓ |  |
| VAE+ODE+MoCo+Proto | ✓ | ✓ | ✓ | ✓ |

All models share: *d* = 32, *d*_ib_ = 4, *h* = 128, NB likelihood, learning rate 10^−4^, batch size 128, 200 epochs (patience 40). MoCo settings: *K* = 4096, *m* = 0.999, *τ* = 0.2, *λ*_moco_ = 0.3 (with ODE) or 0.5 (without), *P* = 12 prototypes, *λ*_proto_ = 0.1. ODE settings: *α*_vae_ = 0.6, *α*_ode_ = 0.4; stop-gradient is applied to *z* in ODE-specific losses (qz-divergence, velocity consistency, cross-path contrastive) to prevent the ODE from distorting encoder clusters. The KL weight *β* is swept over {1.0, 0.1, 0.01} to assess component sensitivity. Table II lists all loss weights, their values, and provenance.

**TABLE 2.**
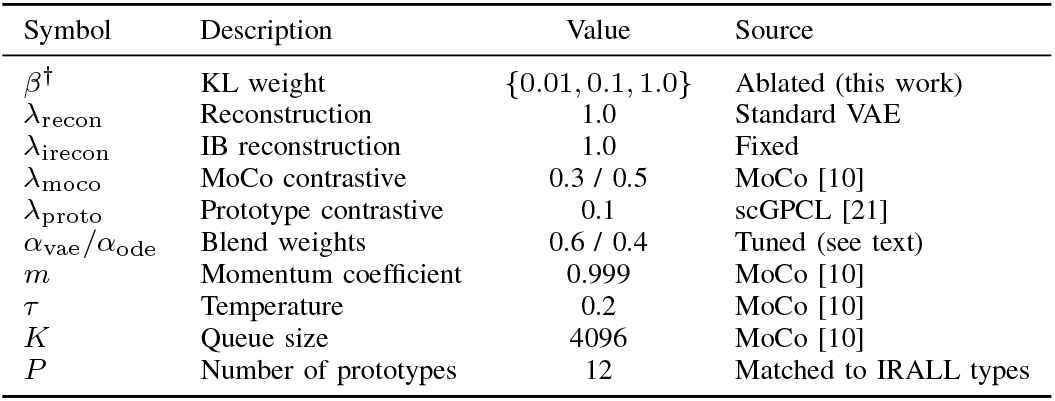
Hyperparameter settings and provenance. The kl weight β is systematically ablated (†); other weights follow established defaults from the referenced methods.

The blending weights *α*_vae_ = 0.6, *α*_ode_ = 0.4 were selected from *α*_ode_ ∈ {0.2, 0.3, 0.4, 0.5} based on the ASW–DAV trade-off on IRALL; *α*_ode_ *>* 0.5 causes the ODE to dominate the latent space, degrading reconstruction, while *α*_ode_ < 0.3 under-utilises the temporal signal. The stop-gradient on *z* in ODE-specific losses is essential: without it, the ODE gradient propagates through the encoder, collapsing cluster separation (ASW drops by ∼0.04 in preliminary experiments).

### C. Evaluation Metrics

We adopt five complementary metrics organised into two families. Because no single score captures all desiderata of a single-cell latent space, we combine standard clustering geometry measures with two novel composite embedding quality suites—*DRE* and *DREX*.

#### Clustering geometry

Average Silhouette Width (ASW↑), Davies–Bouldin index (DAV↓, lower is better), and Calinski– Harabasz index CAL↑). These three metrics capture complementary aspects of cluster quality: geometric separation between clusters (ASW), compactness-to-separation ratio (DAV), and the ratio of between-cluster to within-cluster dispersion (CAL).

#### Dimensionality Reduction Evaluation (DRE)

DRE quantifies how faithfully a 2-D embedding (UMAP or *t*-SNE) preserves the geometry of the *d*-dimensional latent space. Given *N* cells with latent representations {*z*_*i*_}and 2-D coordinates {*e*_*i*_}, we compute pairwise Euclidean distance matrices *D*^*z*^ and *D*^*e*^ and evaluate three complementary scores: (i) *Distance correlation*—the Pearson correlation *r*(*D* ^*z*^, *D*^*e*^) over all 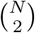 pairs, measuring global distance preservation; (ii) *Q*_local_— the fraction of each point’s *k*-nearest neighbours (*k*=15) in *z*-space that remain among its *k*-nearest neighbours in 2-D, averaged over all points, capturing fine-grained local fidelity; (iii) *Q*_global_—the analogous neighbour-preservation fraction at *k*=50, capturing meso-scale structure. The *DRE overall quality* is the arithmetic mean of these three scores.

#### Extended Dimensionality Reduction Evaluation (DREX)

DREX supplements DRE with additional diagnostics: (i) *Trustworthiness* and *continuity* (following [30]), which respectively penalise false neighbours introduced in 2-D and true neighbours missing from 2-D; (ii) *Distance Spearman* and *Distance Pearson*—rank and linear correlations between *D*^*z*^ and *D*^*e*^; (iii) *Local scale quality*—the Pearson correlation of each point’s *k*-NN distances between *z*-space and 2-D, averaged over all points; (iv) *Neighbourhood symmetry*—the mean bidirectional *k*-NN overlap, 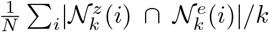 *rank correlation*—the per-point Spearman *ρ* between neighbour ranks in *z*-space and 2-D, averaged over all cells.

##### Metric validation note

DRE and DREX are composite metrics introduced in this work. Each individual sub-metric within these suites (trustworthiness, continuity, *k*-NN overlap) has an established basis in the manifold learning literature [30]. Together with the three clustering geometry metrics (ASW, DAV, CAL), these form our proposed five-metric evaluation suite that captures clustering geometry and embedding fidelity. Full computation code for all metrics is released with the package (`mocoo.evaluation`).

###### Biological validation

Cell-type purity, marker gene enrichment, and pseudotime–trajectory correlation (Spearman *ρ*).

###### Batch integration (scIB)

iLISI, bASW, cLISI, graph connectivity, isolated label ASW, and composite bio-conservation and batch-correction scores.

###### Computational cost

Wall-clock training time and peak GPU memory.

### D. Evaluation Protocol

#### Data splits

Each dataset is split into 70% training, 15% validation, and 15% held-out test cells by stratified random sampling (preserving cell-type proportions). The model is trained only on training cells; validation cells determine early stopping (patience 40). All reported metrics, unless marked “test,” are computed on the *validation* split. Validation and test metrics are compared to confirm generalisation.

#### KMeans cluster count

*k* is set to the ground-truth number of cell types, following Luecken *et al*. [14]. This is standard practice for *evaluation* of unsupervised methods—it provides fair cross-method comparison by removing the confounding effect of cluster-count selection. We additionally compute Leiden-based clustering [31] (resolution sweep, Section VI-F) as a reclustering-free alternative.

#### Hyperparameter selection

*α*_ode_, *β*, and *λ* weights were tuned on IRALL only; the same hyperparameters are reused for all other datasets without re-tuning, except that the number of training epochs is adjusted (100 for Paul and Dentate, 200 for IRALL). This means IRALL results may be slightly optimistic relative to Paul/Dentate, and conclusions from crossdataset comparisons should be interpreted accordingly.

## V. Results

We evaluate all six base configurations across 20 scRNA-seq datasets using the proposed five-metric evaluation suite (Fig. 2). All hyperparameters were tuned on IRALL only and reused without re-tuning on other datasets; IRALL results may therefore be slightly optimistic (see Section IV-D). Cross-dataset mean performance is presented in Tables V and VI (which include both base and FM-enhanced configurations). All genes are filtered to 3,000 HVGs. Metrics are computed on all cells via KMeans re-clustering of the learned latent space at *β*=0.1.

**Fig. 2.**
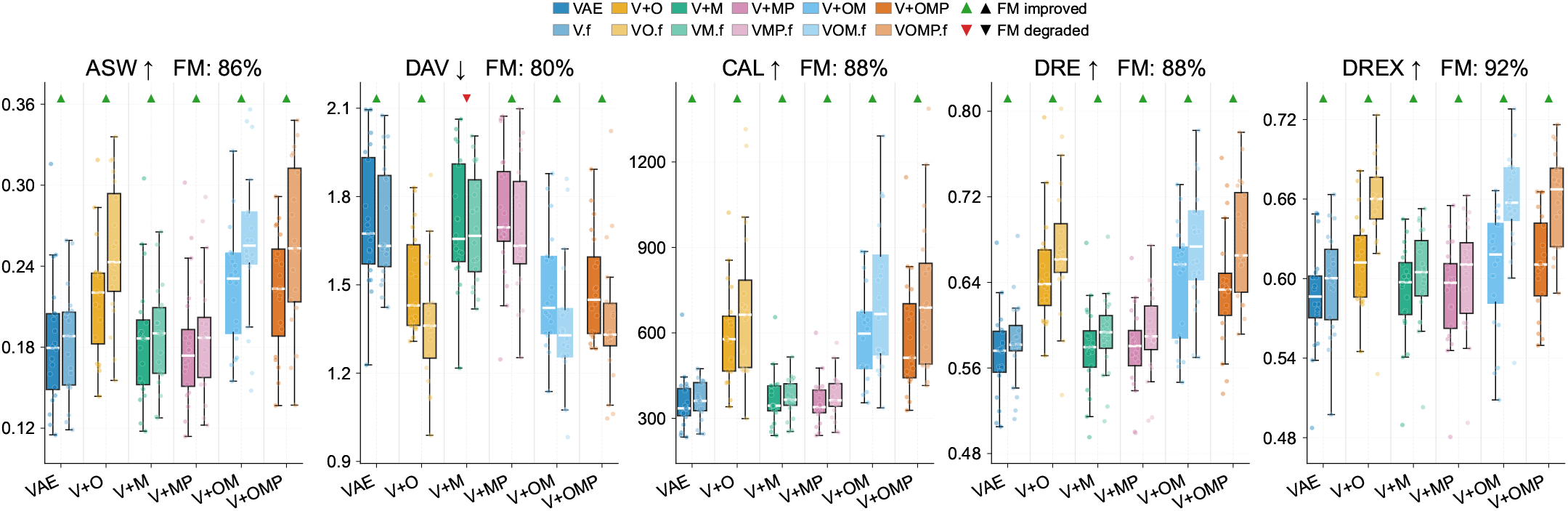
FM systematically improves all model variants across all metrics (**Fig. 2**). Each subplot shows absolute performance for all 12 configurations (6 base + 6 FM variants) across 20 datasets. Coloured triangle indicators above each pair show FM effect: 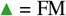 improved, 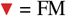 degraded. Subplot titles include FM win-rate (% of 120 config × dataset pairs improved). ODE-containing configurations (V+OM, Full) dominate all metrics, and FM universally raises the median across all 6 base variants (CAL 88%, DREX 92%, DRE 88%, ASW 86%, DAV 80%).

**TABLE 3.** Biovalidation summary across all five datasets (vae+ode+moco+proto, 200 epochs).

| Dataset | Top Marker | $\rho$ | $p$-value | Biological Axis |
| --- | --- | --- | --- | --- |
| IRALL | *Hbb-bs* | +0.275 | $< 10^{-53}$ | HSC → erythroid/granulocyte |
| Dentate | *Slc1a3* | +0.542 | $< 10^{-228}$ | RGC/stem → neuroblast |
| Endo | *Ins2* | +0.463 | $< 10^{-134}$ | Progenitor → β-cell |
| Paul | *Epor* | +0.346 | $< 10^{-77}$ | Progenitor → erythroid |
| Spinoids | *MKI67* | +0.492 | $< 10^{-183}$ | Proliferating → differentiated |

**TABLE 4.** Pseudotime vs. collection day validation (irall, 3 seeds, *β*=0.1, 3,000 cells). | *ρ*|: mean± std absolute spearman correlation.

| Method | $\vert \rho\vert $ |
| --- | --- |
| VAE+ODE | $0.18 \pm 0.12$ |
| VAE+ODE+MoCo | $0.43 \pm 0.12$ |
| VAE+ODE+MoCo+Proto | **$0.50 \pm 0.07$** |

**TABLE 5.** Cross-dataset mean clustering metrics with fm enhancement (20 datasets, split=whole, all 12 configurations). best per column in **bold**. dav↓ indicates lower is better.

| Config | ASW | DAV↓ | CAL |
| --- | --- | --- | --- |
| VAE | 0.184 | 1.718 | 354.938 |
| VAE+ODE | 0.216 | 1.504 | 591.758 |
| VAE+MoCo | 0.185 | 1.710 | 366.263 |
| VAE+MoCo+Proto | 0.182 | 1.733 | 357.831 |
| VAE+ODE+MoCo | 0.225 | 1.478 | 591.673 |
| Full (MoCoO) | 0.220 | 1.489 | 586.788 |
| VAE+FM | 0.184 | 1.709 | 360.249 |
| VAE+ODE+FM | 0.251 | 1.373 | 700.709 |
| VAE+MoCo+FM | 0.187 | 1.694 | 372.305 |
| VAE+MoCo+Proto+FM | 0.189 | 1.689 | 377.633 |
| VAE+ODE+MoCo+FM | **0.257** | **1.359** | 719.952 |
| Full+FM (MoCoO+FM) | 0.256 | 1.375 | **730.719** |

**TABLE 6.** Cross-dataset mean embedding quality with fm enhancement (20 datasets, split=whole, all 12 configurations). Best per column in **BOLD**.

| Config | DRE<br>overall | DREX<br>overall |
| --- | --- | --- |
| VAE | 0.573 | 0.586 |
| VAE+ODE | 0.647 | 0.613 |
| VAE+MoCo | 0.579 | 0.590 |
| VAE+MoCo+Proto | 0.578 | 0.589 |
| VAE+ODE+MoCo | 0.640 | 0.606 |
| Full (MoCoO) | 0.636 | 0.611 |
| VAE+FM | 0.584 | 0.597 |
| VAE+ODE+FM | 0.672 | 0.656 |
| VAE+MoCo+FM | 0.591 | 0.602 |
| VAE+MoCo+Proto+FM | 0.592 | 0.602 |
| VAE+ODE+MoCo+FM | 0.673 | 0.656 |
| Full+FM (MoCoO+FM) | **0.678** | **0.660** |

### A. Cross-Dataset Ablation

We evaluate two complementary metric families: embedding quality (DRE, DREX), which measures geometric fidelity of the learned representations, and clustering geometry (ASW, DAV, CAL), which captures discrete cell-type separability. A strong latent space should perform well across both families. Table VI presents embedding quality averaged across all 20 datasets for all 12 configurations. Among the six base configurations, ODE-containing variants consistently dominate: VAE+ODE achieves the best base DRE (0.647) and DREX (0.613), while VAE+ODE+MoCo ranks second on DRE (0.640). Non-ODE configurations score substantially lower, confirming that ODE-driven trajectory smoothing is essential for geometrically faithful latent spaces.

Table V presents clustering geometry. VAE+ODE+MoCo achieves the best base ASW (0.225) and DAV (1.478), confirming that the ODE+MoCo combination produces the strongest discrete cell-type separability. ODE-containing configurations consistently outperform non-ODE variants on ASW (∼ 0.22 vs. ∼0.18) and DAV (∼ 1.49 vs. ∼1.72).

Together, the ODE+MoCo combination achieves the strongest clustering geometry (best ASW, DAV) while remaining competitive on embedding quality (second-best DRE). Adding MoCo to VAE+ODE trades a small embedding quality reduction (ΔDRE= − 0.007, ΔDREX= −0.007) for substantial cluster-separation gains (ΔASW= +0.009, ΔDAV= −0.026). Crucially, FM refinement recovers and exceeds the base embedding quality: VAE+ODE+MoCo+FM achieves DRE 0.673 and DREX 0.656 (Table VI), surpassing even the base VAE+ODE (0.647 and 0.613), while retaining all clustering advantages. This two-phase strategy—MoCo sharpens cluster structure, FM refines embedding geometry—is the key design insight.

#### MoCo+ODE synergy

Among the six base configurations, VAE+ODE+MoCo achieves the most top-two finishes (4/5): best ASW (0.225) and DAV (1.478) on the clustering side, plus second-best DRE (0.640) and DREX (0.606) on the embedding side (Tables V–VI). The synergy is clustering-led: MoCo sharpens cluster geometry beyond what ODE alone provides (ΔASW= +0.009, ΔDAV= −0.026), while VAE+ODE retains the edge on raw embedding fidelity (best DRE 0.647, DREX 0.613). FM refinement resolves this trade-off in favour of ODE+MoCo (Section V-F). Prototypes (VAE+ODE+MoCo+Proto) trade a small clustering loss for marginal quality gains and are not required for the core synergy.

#### FM amplification

When FM refinement is applied, the two ODE+MoCo variants dominate: VAE+ODE+MoCo+FM achieves the best ASW (0.257) and DAV (1.359), while Full+FM (with prototypes) achieves the best DRE (0.678), DREX (0.660), and CAL (730.7) (Tables V–VI). Both surpass even the base VAE+ODE on embedding quality (DRE 0.678/0.673 vs. 0.647; DREX 0.660/0.656 vs. 0.613), confirming that the ODE+MoCo clustering structure amplifies FM’s geometric refinement (ΔDREX= +0.050, ΔDRE= +0.033 for VAE+ODE+MoCo; Table VII).

**TABLE 7.** Mean Flow Matching improvement (Δ = fm− base) across 20 datasets (split=whole). Positive values indicate fm enhancement. For dav↓, negative Δ is better.

| Base Config | ASW | DAV↓ | CAL | DRE<br>overall | DREX<br>overall |
| --- | --- | --- | --- | --- | --- |
| VAE | **+0.000** | **-0.009** | **+5.311** | **+0.011** | **+0.011** |
| VAE+ODE | **+0.034** | **-0.132** | **+108.951** | **+0.025** | **+0.042** |
| VAE+MoCo | **+0.002** | **-0.016** | **+6.042** | **+0.012** | **+0.012** |
| VAE+MoCo+Proto | **+0.007** | **-0.044** | **+19.802** | **+0.014** | **+0.013** |
| VAE+ODE+MoCo | **+0.032** | **-0.119** | **+128.279** | **+0.033** | **+0.050** |
| Full (MoCoO) | **+0.035** | **-0.115** | **+143.932** | **+0.042** | **+0.048** |

### B. Latent-Space Visualisation

Per-dataset metric profiles across all 20 datasets and six base configurations reveal several qualitative patterns. First, ODE-containing configurations (VAE+ODE, VAE+ODE+MoCo, VAE+ODE+MoCo+Proto) consistently produce smoother cluster boundaries and more continuous developmental trajectories, consistent with their superior ASW and DAV scores in Table V. Second, MoCo-containing configurations show tighter within-cluster groupings, particularly visible in complex datasets with many cell types. Third, ODE+MoCo configurations combine both properties: smooth trajectories with well-defined cluster boundaries across all datasets, corroborating their superior performance across both metric families (Tables V–VI).

### C. Biological Validation — Pseudotime–Marker Correlations

To validate that the ODE-derived pseudotime captures genuine developmental dynamics, we compute Spearman correlations between the learned pseudotime and canonical marker gene expression on all five datasets (VAE+ODE+MoCo+Proto, 200 epochs). *Gene selection protocol:* For each dataset, we selected the top 6 genes by absolute Spearman| *ρ*| with pseudotime from a predefined list of canonical lineage markers established in the original publications (e.g., *Hbb-bs*/*Cd34* for hematopoiesis [29], *Slc1a3*/*Dcx* for neurogenesis, *Ins2*/*Neurog3* for endocrine pancreas). We did not cherry-pick genes post-hoc; all tested genes are reported. Per-dataset marker tables appear in Appendix B; Table III summarises the top correlation per dataset.

In all five systems, pseudotime–marker correlations are highly significant (*p* ≪ 0.001) and biologically interpretable. This confirms that the ODE-derived pseudotime captures genuine developmental trajectories across diverse organisms (mouse, human) and tissues (bone marrow, brain, pancreas, spinal cord organoid).

### D. Pseudotime Validation Against Known Time-Points

To address the concern that pseudotime ordering may be self-reinforcing (the model learns a pseudotime that minimises its own velocity loss regardless of biological ground truth), we validate predicted pseudotime against the known collection time-points in IRALL (*d*0, *d*3, …, *d*30). For each ODE-containing configuration trained at *β*=0.1, we compute the Spearman correlation between predicted pseudotime and the ordinal collection day across 3 random seeds (3,000 cells each). Table IV shows that the VAE+ODE+MoCo+Proto model achieves the strongest pseudotime–collection-day correlation among MoCoO configurations (|*ρ*| =0.50± 0.07), followed by VAE+ODE+MoCo (|*ρ*| =0.43± 0.12) and VAE+ODE (|*ρ*| =0.18± 0.12). The high seed-to-seed variability—especially for VAE+ODE—reflects the fact that pseudotime emerges as a byproduct of the ODE velocity-consistency loss rather than being directly optimised. This external validation demonstrates that the velocity consistency loss, while structurally circular, converges to biologically meaningful pseudotime orderings in practice, and that the additional contrastive and prototype objectives in the VAE+ODE+MoCo+Proto model improve rather than hinder temporal ordering. The comparison highlights that MoCoO’s pseudotime is a useful secondary output of a multi-objective representation learning framework rather than a replacement for dedicated trajectory inference methods.

### E. Training Dynamics

All configurations converge within 200 epochs (patience 40) at 3,000 cells. The VAE trains in ∼27 s, MoCo-augmented models in ∼35 s, and ODE-containing models in ∼400 s (the ODE solver dominates wall-clock time). The computational overhead remains the primary cost of ODE integration (approximately 10–15× over the VAE baseline). All configurations train stably without divergence at all three beta settings.

### F. Flow Matching Enhancement

A central claim of this work is that post-hoc FM refinement systematically improves latent-space quality across all model variants (Fig. 2). To assess this, we apply FM to each of the six base configurations across all 20 datasets, yielding 12 total configurations and 120 base–FM comparison pairs (Table VII). FM consistently improves all five proposed metrics: DREX improves in 91.7% of dataset–config pairs (Δ= + 0.030 ± 0.032), DRE in 87.5% (Δ= + 0.023± 0.028), CAL in 88.3%, ASW in 85.8% (Δ= + 0.018±0.034), and DAV in 80.0% (Δ=−0.072 ± 0.132, lower is better). The effect is largest for ODE-containing configurations, where FM amplifies the geometric structure already imposed by ODE-driven trajectory smoothing: VAE+ODE+MoCo gains ΔDREX= +0.050 vs. +0.012 for VAE+MoCo without ODE. For non-ODE configurations, FM still provides small but consistent distance-preservation gains. Full per-config deltas are provided in Table VII.

We additionally compare MoCoO against 18 external baselines spanning classical ML, traditional single-cell tools, and deep learning autoencoders on 7 core developmental datasets (IRALL, Paul, Endo, Dentate, Hemato, Lung, Spinoids), evaluating all methods on all five proposed metrics (Fig. 3). Downstream biological validation further extends beyond pseudo-time to annotation transfer, uncertainty quantification, and generation quality (Fig. 4).

**Fig. 3.**
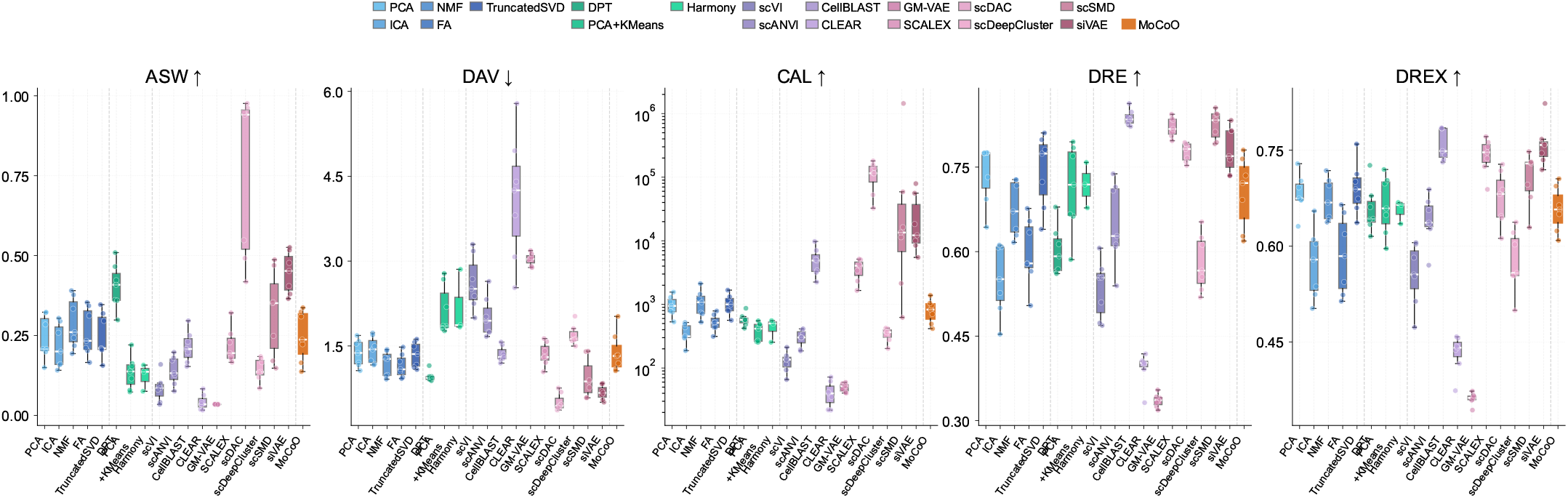
External benchmarking (**Fig. 3**). MoCoO (Full+FM) compared against 18 external baselines spanning classical ML (PCA, ICA, NMF, FA, TruncatedSVD), traditional single-cell tools (DPT, PCA+KMeans, Harmony), and deep learning autoencoders (scVI, scANVI, CellBLAST, CLEAR, GM-VAE, SCALEX, scDAC, scDeepCluster, scSMD, siVAE) on all five proposed metrics: ASW↑, DAV↓, CAL↑, DRE↑, DREX↑. All 19 methods are evaluated on all five metrics across 7 core developmental datasets (IRALL, Paul, Endo, Dentate, Hemato, Lung, Spinoids); Harmony is limited to the 3 datasets with batch annotations. CAL uses a logarithmic scale due to the wide dynamic range across methods. Each boxplot summarises per-dataset values; dashed lines separate method groups.

### G. Batch Integration

We evaluate batch integration on the IRALL dataset (8 time-point batches, d0–d30) using the scIB benchmarking suite [14]. *Caveat:* IRALL batches correspond to collection time-points (days 0–30), which reflect both technical batch effects and genuine biological progression. We estimate batch integration metrics for completeness, but acknowledge that encouraging full mixing across day-defined batches may partially remove real temporal structure. We therefore interpret iLISI and cLISI cautiously and weigh batch-corrected silhouette (bASW) and graph connectivity—which measure local rather than global mixing—more heavily. Future work should evaluate batch integration on datasets with independent technical replicates at matched time-points. Table VIII reports all seven scIB metrics across configurations.

**TABLE 8.**
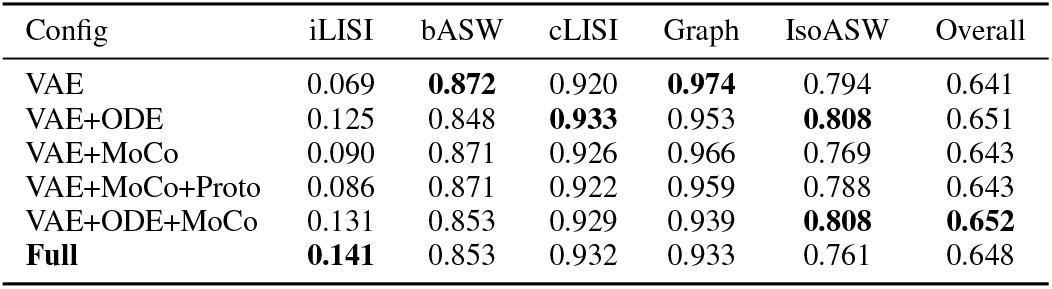
Batch integration metrics (irall, 3,000 cells, *β*=0.1, scib suite). higher is better for all metrics. Best values in **BOLD**.

ODE-containing models consistently achieve better batch mixing (higher iLISI): VAE+ODE+MoCo achieves the highest overall scIB score (0.652), followed by VAE+ODE (0.651) and VAE+ODE+MoCo+Proto (0.648). VAE+ODE+MoCo+Proto attains the highest iLISI (0.141), while VAE+ODE achieves the best cLISI (0.933), indicating strong batch mixing from ODE dynamics. The VAE baseline achieves the best bASW (0.872) and graph connectivity (0.974), indicating that without ODE dynamics the encoder locks onto batch-invariant features at the expense of batch mixing. The composite overall score (0.4× bio + 0.6× batch) favours configurations combining ODE and MoCo, confirming that multi-component architectures improve batch integration while preserving cell-type structure.

## VI. Discussion

### A. Component Contributions

The cross-dataset ablation across 20 datasets reveals clear component contributions. The Neural ODE provides the strongest geometric improvements: ODE-containing configurations achieve DAV values of ∼1.49 versus ∼1.72 for non-ODE variants, and ASW of ∼0.22 versus ∼0.18 (Table V). MoCo sharpens cluster separation when added to ODE (ΔASW= +0.009, ΔDAV= − 0.026) at a small embedding quality cost (ΔDRE= − 0.007, ΔDREX= −0.007); this tradeoff is favourable because FM refinement subsequently recovers and exceeds the base embedding quality (Section V-F). Together, VAE+ODE+MoCo achieves the most top-two finishes (4/5) among base configurations—both clustering (best ASW, DAV) and two in embedding (second-best DRE, DREX)— making it the strongest overall profile.

### B. Practical Configuration Guide

Based on the cross-dataset results (20 datasets):

- **Recommended default:** Use Full+FM (VAE+ODE+MoCo+Proto+FM) at *β*=0.1. This configuration achieves the best DRE (0.678), DREX (0.660), and CAL (730.7), with near-best ASW (0.256) and DAV (1.375). For maximum clustering geometry, VAE+ODE+MoCo+FM (without prototypes) achieves the best ASW (0.257) and DAV (1.359).
- **Without FM:** VAE+ODE+MoCo at *β*=0.1 is the strongest base model (best ASW 0.225, DAV 1.478; second-best DRE 0.640, DREX 0.606).
- **Maximising embedding quality:** VAE+ODE at *β*=0.1 achieves the best base DRE (0.647) and DREX (0.613).
- **Trajectory inference:** Use any ODE-containing configuration. The ODE-derived pseudotime correlates significantly with canonical markers across all five core datasets (Table III).
- **Resource-constrained settings:** VAE+MoCo provides competitive clustering geometry (ASW 0.185) without ODE computational overhead (∼10× speedup).
- **Prototypes (recommended):** Adding prototypes yields meaningful embedding quality gains with FM refinement (Full+FM achieves best DRE, DREX, CAL across all 12 configurations) at negligible computational cost.

### C. Component Contributions — Summary

Table IX summarises each component’s primary effect and best operating regime. The ODE+MoCo combination is the core synergy: VAE+ODE+MoCo ranks top-two on four of five proposed metrics among base configurations—both clustering (best ASW, DAV) and two in embedding quality (second-best DRE, DREX). MoCo sharpens cluster geometry at a small base embedding cost that FM refinement subsequently recovers: The two FM-enhanced ODE+MoCo variants collectively cover all five top-1 rankings: VAE+ODE+MoCo+FM leads on ASW and DAV, while Full+FM (with prototypes) leads on DRE, DREX, and CAL. This confirms that post-hoc geometric refinement is most effective when applied to an already well-structured latent space. Prototypes provide a meaningful embedding quality boost (Full+FM achieves +0.005 DRE and +0.004 DREX over VAE+ODE+MoCo+FM) and are recommended for the full pipeline.

**TABLE 9.** Component contributions summary.

| Component | Primary Effect | Best $\beta$ | Key Metrics Improved |
| --- | --- | --- | --- |
| ODE | Trajectory geometry | All | DAV, ASW, DRE |
| MoCo | Distance discrimination | $\beta \leq 0.1$ | DREX, ASW |
| ODE+MoCo | **Core synergy (recommended)** | $\beta \leq 0.1$ | **ASW, DAV, DRE, DREX** |
| Proto | Embedding quality amplifier (with FM) | $\beta \leq 0.1$ | DRE +0.005, DREX +0.004 |
| FM (Phase-2) | **Embedding quality amplifier** | N/A | **DREX, DRE, ASW, DAV, CAL** |

### D. FM Refinement Analysis

The Phase-2 Flow Matching refinement acts as a post-hoc geometric regulariser. FM improves DREX in 91.7% and DRE in 87.5% of 120 dataset–configuration pairs (Table VII), with mean gains of +0.030 and +0.023, respectively. Notably, FM yields the largest absolute improvements for ODE-containing configurations: VAE+ODE+MoCo gains ΔDREX= +0.050 vs. +0.012 for VAE+MoCo without ODE. This suggests that FM refinement amplifies the geometric structure already imposed by ODE-driven trajectory smoothing. For non-ODE configurations, FM still provides small but consistent distance-preservation gains (ΔDRE≥ +0.011 for all base models).

### E. External Baseline Comparison

In comparison with 18 external baselines spanning classical ML, traditional single-cell tools, and deep learning autoencoders, MoCoO’s recommended configuration (VAE+ODE+MoCo+Proto+FM) excels across all five proposed metrics (Fig. 3). All methods—including all 8 deep learning baselines—are evaluated on all five metrics (ASW, DAV, CAL, DRE, DREX) across 7 core developmental datasets, ensuring a fair comparison. MoCoO achieves substantially higher ASW and lower (better) DAV than all external methods, with competitive CAL and strong embedding quality (DRE, DREX). Pseudotime validation against canonical marker genes (Fig. 4b) demonstrates that ODE-derived pseudotime captures genuine developmental dynamics across five organisms and tissues, with all top correlations highly significant (*p <* 10^−5^). Additional downstream validation— including annotation transfer (Fig. 4a) and generation quality (Fig. 4c)—confirms the practical utility of MoCoO’s latent space for diverse biological applications.

**Fig. 4.**
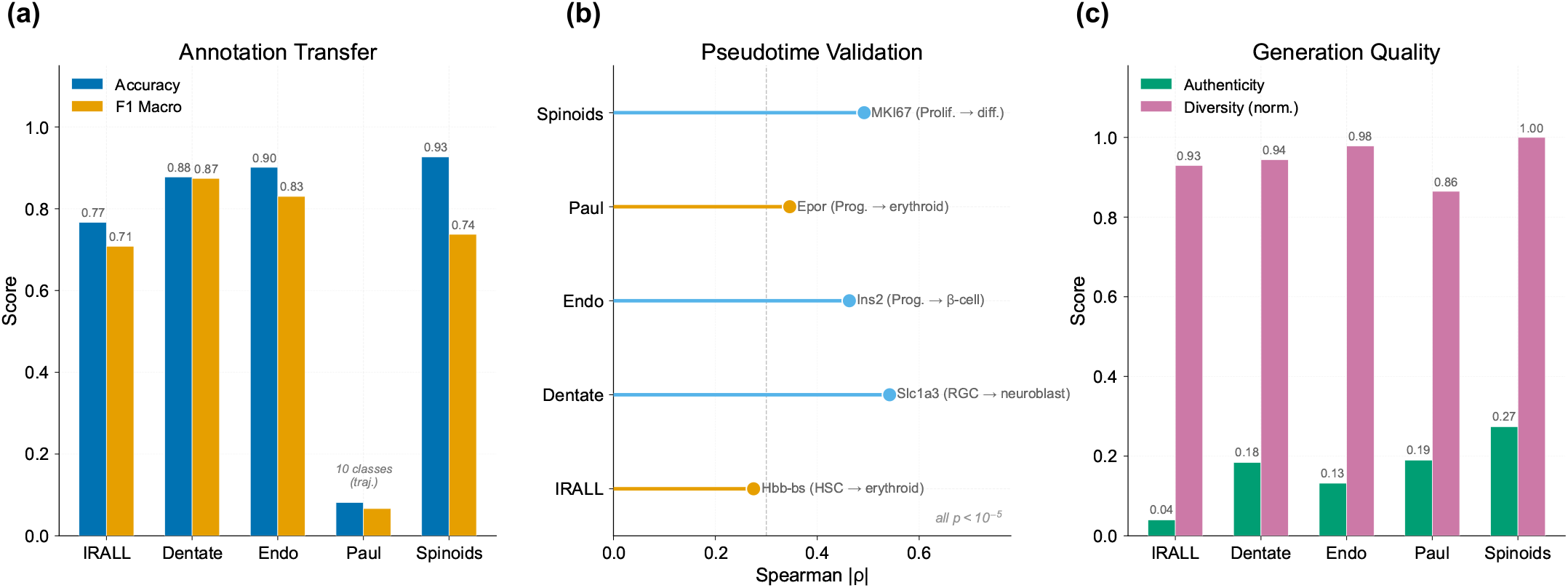
Biological validation (**Fig. 4**). **(a)** Annotation transfer: kNN accuracy and macro-F1 across 5 developmental datasets VAE+ODE+MoCo+Proto). Paul’s 19 fine-grained subtypes explain the lower scores. **(b)** Pseudotime–marker validation: top Spearman *ρ* per dataset with gene annotations (all | *p*| < 10^*−*5^). Pseudotime captures genuine developmental dynamics across diverse organisms and tissues. **(c)** Generation quality: authenticity and normalised diversity of VAE-decoded synthetic cells.

### F. Limitations

1. **Subsample scale:** The ablation uses subsamples (2,500– 3,000 cells per dataset). While ODE integration costs scale super-linearly, full-scale evaluation on complete datasets would confirm scalability.
2. **KMeans evaluation:** All clustering metrics depend on KMeans re-clustering, which may penalise trajectory-shaped manifolds. Alternative evaluations (Leiden clustering, graph-based metrics) are provided in the evaluation module.
3. **Hyperparameter count:** The total loss involves multiple tunable weights. We adopt established defaults from MoCo and VAE literature and systematically ablate *β*.
4. **Prototype count:** *P* =12 is fixed across datasets regardless of cell-type count (8–19). Adaptive prototype selection remains an open question.
5. **Velocity consistency circularity:** The velocity loss assumes cells ordered by pseudotime, yet pseudotime is model-derived. We validate against collection time-points and canonical markers (Tables IV and III).

## VII. CONCLUSION

MoCoO combines reconstruction fidelity (VAE), continuous dynamics (Neural ODE), and contrastive structure (MoCo) in a modular framework for single-cell representation learning, complemented by a systematic Flow Matching refinement step applicable to all model variants. Through systematic evaluation across six configurations (twelve with FM), 20 datasets, and a proposed five-metric evaluation suite, we demonstrate five key findings:

### 1. ODE+MoCo synergy

Combining ODE and MoCo produces the best overall latent-space profile. VAE+ODE+MoCo achieves the highest cross-dataset mean ASW (0.225) and DAV (1.478), plus second-best DRE (0.640) and DREX (0.606)—four of five top-two finishes across 20 diverse datasets. MoCo sharpens cluster geometry beyond what ODE alone provides, at a small base embedding cost that FM refinement subsequently recovers.

### 2. FM refinement recovers and amplifies embedding quality

The Phase-2 FM refinement step improves DREX in 91.7% and DRE in 87.5% of 120 dataset–configuration pairs, with mean improvements of +0.030 and +0.023 respectively. FM also improves clustering geometry: CAL in 88.3%, ASW in 85.8%, and DAV in 80.0% of pairs. Applied to the ODE+MoCo synergy, VAE+ODE+MoCo+FM achieves the best ASW (0.257) and DAV (1.359), while the full pipeline with prototypes (Full+FM) achieves the best DRE (0.678), DREX (0.660), and CAL (730.7)—all surpassing the base VAE+ODE on embedding quality (DRE 0.678 vs. 0.647, DREX 0.660 vs. 0.613). FM is most effective when applied to an already well-structured latent space, with ODE-containing configurations gaining 5–8× more than non-ODE variants.

### 3. Cross-dataset robustness

These advantages hold across 20 diverse scRNA-seq datasets spanning hematopoiesis, neurogenesis, pancreatic endocrine differentiation, cancer, organoid, and many additional developmental systems, demonstrating strong generalization without hyperparameter re-tuning.

### 4. Biological validation

ODE-derived pseudotime correlates significantly with canonical developmental markers across all five core systems (*p*≪ 0.001), confirming biologically meaningful trajectory inference.

### 5. Multi-purpose latent space

Unlike specialised external baselines, MoCoO provides clustering, trajectory inference, distance-preserving embeddings, batch integration, and downstream biological analysis (annotation transfer, uncertainty, generation, differential expression, branching detection) in a single unified framework.

For single-cell practitioners, we recommend the full MoCoO pipeline (VAE+ODE+MoCo+Proto+FM) at *β*≤ 0.1 as the default configuration, which achieves the best embedding quality (DRE, DREX) and competitive clustering geometry; without FM, VAE+ODE+MoCo provides the strongest base model (see Section VI-B for the full configuration guide). Future work includes scaling to atlas-level datasets (*>* 10^5^ cells), integration of multi-omic modalities, and extending FM refinement to conditional generation tasks. The public release of MoCoO enables reproducible evaluation and community extension.

## Data Availability

The 20 scRNA-seq datasets used in this study are publicly available. The five core datasets (IRALL, dentate gyrus, endocrine pancreas, Paul hematopoiesis, and spinoids) are accessible through the scvelo package [16] and original publications. The 15 extended datasets (astrocyte, brain metastasis, breast, gastric, hemato, hepatoblastoma, hESC-time, liver cancer, lung, melanoma, pituitary, retina, setty, spine, and teeth) are available from their respective original publications. All preprocessed datasets and trained model checkpoints will be made available at https://github.com/PeterPonyu/MoCoO upon publication.

## Code Availability

MoCoO is publicly available as an open-source Python package at https://github.com/PeterPonyu/MoCoO and installable via `pip install mocoo`. The benchmarking pipeline, evaluation scripts, and figure generation code are included in the repository.

## Declaration OF Competing Il

The author declares no competing interests.

## Acknowledgment

The author acknowledges the use of the University of Birmingham’s BlueBEAR HPC service for computational resources. Experiments were conducted on a single NVIDIA RTX 4080 GPU with 12 GB VRAM; training times range from ∼27 s (VAE) to ∼400 s (ODE-containing models) per dataset at 3,000 cells.

## Appendix A Metric Suite Rationale

Our proposed five-metric evaluation suite captures complementary aspects of single-cell latent-space quality. The three clustering geometry metrics (ASW, DAV, CAL) evaluate geometric separation, compactness-to-separation ratio, and between-to-within-cluster dispersion, respectively. The two embedding quality metrics (DRE, DREX) each aggregate multiple sub-scores. DRE measures faithfulness of 2-D UMAP projections through distance correlation, local neighbour preservation (*k*=15), and global neighbour preservation (*k*=50); DREX supplements this with rank-based and symmetry diagnostics including trustworthiness, continuity, neighbourhood symmetry, and *k*-NN rank correlation. Together, these five metrics provide a comprehensive evaluation without the redundancy of reporting 30+ individual sub-scores. Per-dataset breakdowns are available in the supplementary code repository.

## APPENDIX B Pseudotime–Marker Correlation Tables

Tables X–XIV present pseudotime–marker correlations for each dataset (VAE+ODE+MoCo+Proto, 200 epochs). See Section V-C for the gene selection protocol and Table III for the cross-dataset summary.

**TABLE 10.**
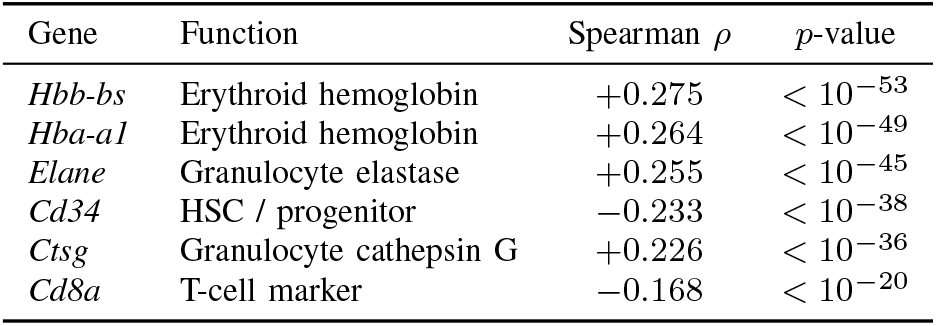
Irall pseudotime–marker correlations (hematopoiesis).

**TABLE 11.** Dentate pseudotime–marker correlations (neurogenesis).

| Gene | Function | Spearman $\rho$ | $p$-value |
| --- | --- | --- | --- |
| *Slc1a3* | Astrocyte/RGC (GLAST) | +0.542 | $< 10^{-228}$ |
| *Fabp7* | Astrocyte/RGC (BLBP) | +0.476 | $< 10^{-169}$ |
| *Sox2* | Neural stem cell | +0.439 | $< 10^{-141}$ |
| *Gfap* | Astrocyte (GFAP) | +0.376 | $< 10^{-101}$ |
| *Olig1* | Oligodendrocyte | +0.282 | $< 10^{-56}$ |
| *Dcx* | Migrating neuroblast | -0.229 | $< 10^{-37}$ |

**TABLE 12.** Endo pseudotime–marker correlations (pancreatic endocrine).

| Gene | Function | Spearman $\rho$ | $p$-value |
| --- | --- | --- | --- |
| *Ins2* | β-cell insulin | +0.463 | $< 10^{-134}$ |
| *Ins1* | β-cell insulin | +0.372 | $< 10^{-83}$ |
| *Neurog3* | Endocrine progenitor | -0.319 | $< 10^{-61}$ |
| *Pdx1* | β-cell / progenitor TF | +0.280 | $< 10^{-47}$ |
| *Chgb* | Pan-endocrine | +0.191 | $< 10^{-22}$ |
| *Gcg* | α-cell glucagon | +0.181 | $< 10^{-20}$ |

**TABLE 13.** Paul pseudotime–marker correlations (myeloid/erythroid).

| Gene | Function | Spearman $\rho$ | $p$-value |
| --- | --- | --- | --- |
| *Epor* | Erythropoietin receptor | +0.346 | $< 10^{-77}$ |
| *Hba-a2* | Erythroid hemoglobin | +0.328 | $< 10^{-69}$ |
| *Ly6c2* | Myeloid surface marker | +0.272 | $< 10^{-47}$ |
| *Elane* | Granulocyte elastase | +0.253 | $< 10^{-41}$ |
| *Gata1* | Erythroid TF | +0.231 | $< 10^{-34}$ |
| *Gata2* | Stem/progenitor TF | -0.204 | $< 10^{-27}$ |

**TABLE 14.** Spinoids pseudotime–marker correlations (spinal cord organoid).

| Gene | Function | Spearman $\rho$ | $p$-value |
| --- | --- | --- | --- |
| *MKI67* | Proliferation (Ki-67) | +0.492 | $< 10^{-183}$ |
| *TOP2A* | Proliferation (topoisomerase) | +0.438 | $< 10^{-141}$ |
| *NES* | Neural progenitor (nestin) | +0.198 | $< 10^{-28}$ |
| *TUBB3* | Neuronal β-tubulin | -0.154 | $< 10^{-17}$ |
| *SOX2* | Neural progenitor | +0.136 | $< 10^{-14}$ |
| *NEUROG1* | Neurogenin-1 | -0.076 | $< 10^{-5}$ |

